# PathoGD: an integrative genomics approach for CRISPR-based target design of rapid pathogen diagnostics

**DOI:** 10.1101/2024.05.14.593882

**Authors:** Soo Jen Low, Matthew O’Neill, William J. Kerry, Natasha Wild, Marcelina Krysiak, Yi Nong, Francesca Azzato, Eileen Hor, Lewis Williams, George Taiaroa, Eike Steinig, Shivani Pasricha, Deborah A. Williamson

**Author notes:** Contributed equally (Shivani Pasricha, Deborah A. Williamson).

## Abstract

The design of highly specific primers and guide RNAs (gRNA) for CRISPR-based diagnostics is often a laborious process. Several tools exist for gRNA design, but most are tailored for genome editing applications. Here, we present PathoGD, an end-to-end bioinformatic pipeline comprising pangenome and *k*-mer modules for rapid and high-throughput design of primers and gRNAs for CRISPR-Cas12a-based pathogen detection. We validated and demonstrated high specificity of a subset of PathoGD-designed primers and gRNAs for the detection of *Neisseria gonorrhoeae* and *Streptococcus pyogenes.* PathoGD will serve as an important resource for designing CRISPR-based diagnostic assays for current and emerging pathogens.

## BACKGROUND

Rapid and accurate diagnosis is essential for the prevention and control of infectious diseases. However, traditional methods such as polymerase chain reaction (PCR) are generally restricted to the laboratory setting due to their reliance on trained personnel and expensive equipment (1). Accordingly, there is a critical need for affordable and accessible point-of-care (POC) diagnostics to increase testing uptake (2). One approach that has gained momentum in recent years leverages clustered regularly interspaced short palindromic repeats (CRISPR) technology. CRISPR technology has found significant applications in infectious disease diagnostics owing to its simplicity, high specificity, and non-reliance on sophisticated equipment (3). CRISPR-based diagnostics have been used for the detection of viruses, bacteria and parasites, including Zika and Dengue virus (4, 5), human papillomavirus (6), *Mycobacterium tuberculosis* (7), *Plasmodium falciparum* (8, 9) and more recently, SARS-CoV-2 (10–12) and *Monkeypox virus* (13, 14). To achieve pathogen detection at low concentrations, CRISPR is often coupled with an upstream isothermal amplification step such as recombinase polymerase amplification (RPA) or loop-mediated isothermal amplification (LAMP).

CRISPR-based diagnostic assays involve a multi-stage process that exploits the unique properties of Cas12 and Cas13 nucleases. The Cas nucleases are directed to bind their target regions by a guide RNA (gRNA) specifically designed to complement the target sequence of interest. In addition to cleaving its target DNA (Cas12) or RNA (Cas13), the Cas nucleases also non-specifically cleave single-stranded DNA (ssDNA) and single-stranded RNA (ssRNA) through target-activated collateral cleavage activity (6, 15–17). Labelling ssDNA and ssRNA molecules with fluorophore-quencher pairs allows detection of target nucleic acids through the fluorescent signal generated by cleavage of reporter molecules. The versatility of the CRISPR-Cas system lies in its ability to be tailored for detection of nucleic acids from virtually any pathogen through design of specific gRNAs that complement the target region.

Crucial to the success of CRISPR-based diagnostics is the selection of a diagnostic target site, followed by design of primers and gRNAs highly specific to the target. Identification of reliable targets and subsequent assay design can be labour-intensive, and typically involves careful analysis of genomic sequences, assessment of conserved regions, consideration of off-target effects, and iterative refinement to achieve optimal performance. Consequently, conventional diagnostic assays often rely on a limited set of marker genes as target regions for amplification, established prior to the next-generation sequencing era, such as *porA* in *Neisseria gonorrhoeae* (18) and *nuc* in *Staphylococcus aureus* (19). This standard approach may fail to capture the full genetic diversity of a pathogen, potentially resulting in diagnostic escape that is only detectable when using a different diagnostic platform (20). CRISPR-Cas12a-based assays also require the presence of a protospacer adjacent motif (PAM), TTTN directly upstream of the gRNA sequence for recognition and DNA binding (21), potentially limiting the use of some well-established diagnostic markers.

Several tools for gRNA design have been developed and are available through a web interface or as a command-line tool. However, most of these were tailored for genome editing applications, primarily in eukaryotic organisms using Cas9. Additionally, some are restricted to a single genome or certain model organisms or contain pre-defined databases, thus limiting their utility, especially for bacterial pathogens. These tools include, but are not limited to, DeepCRISPR (22), GuideMaker (23), CRISPRscan (24), CRISPR-DT (25), CRISPRseek (26), GuideScan (27) and crisprVerse (28). The gRNA design principles for genome editing applications differ from those for CRISPR-based nucleic acid detection or diagnostic applications in multiple ways. In genome editing, the gRNA should bind to a single target site in the genome, whereas multiple gRNA target binding sites in a genome is often advantageous for diagnostic applications. Off-target effects in genome editing are evaluated within the same genome to predict mutations at unintended genomic loci, whereas genomes from different species are often used for evaluating gRNA specificity in diagnostics. Two tools developed specifically for the design of Cas12a gRNAs in diagnostics are PrimedSherlock (29) and CriSNPr (30); however, the former requires primers designed using PrimedRPA (31) or alternative methods as input, while the latter was developed specifically for variant detection. Consequently, there is a need for a fully automated primer and gRNA design tool specific for nucleic acid detection that is scalable and requires minimal user intervention to help researchers overcome the initial hurdle in assay development.

Advancements in computational tools and the availability of pathogen genomic data provide an opportunity to identify alternative diagnostic markers from previously unexplored genomic regions. In this study, we present PathoGD, a bioinformatic pipeline developed to facilitate high-throughput and rapid design of highly specific RPA primers and gRNAs for CRISPR-Cas12a-based pathogen detection. To the best of our knowledge, this is the first bioinformatic tool providing a streamlined end-to-end workflow for primer and gRNA design in pathogens, thereby simplifying the process which normally requires multiple stages. PathoGD leverages available pathogen genomic data to accelerate the *in silico* aspect of CRISPR-Cas12a-based assay design and is scalable to large datasets. We demonstrate the application of PathoGD in designing highly specific primer and gRNAs for five clinically relevant bacterial pathogens and provide experimental validation of these designs for two common bacterial pathogens, *N. gonorrhoeae* and *Streptococcus pyogenes*. In addition, we compare the gRNA designs from PathoGD and PrimedSherlock for six viruses based on a common database of target and non-target genomes.

## RESULTS

### PathoGD pipeline overview

PathoGD is an automated, command-line tool written in bash and R for primer and CRISPR-Cas12a gRNA design that integrates the functionalities of several widely used bioinformatic tools **(Table S1).** PathoGD incorporates two commonly used approaches in comparative genomics, pangenome and *k*-mer analyses (**Figure S1**), either of which can be invoked based on user preferences. These approaches make up the two independent modules of the pipeline, with multiple workflows available under each module. The pipeline takes as input the module and workflow selection, and a configuration file specifying multiple parameters, including: (i) target and non-target taxa; (ii) the National Center for Biotechnology (NCBI) database; (iii) genome assembly level; (iv) gRNA length, and (v) the desired threshold of gRNA prevalence across target genomes. Depending on the module and workflow selected, some of these parameters will be mandatory while others are optional. Users can opt to download genomes automatically from the NCBI Assembly database (32), or alternatively provide their own set of curated target and/or non-target genomes. Genome subsampling can optionally be selected and is particularly useful in reducing computational runtime for certain target species which are overrepresented in the database.

### PathoGD pangenome and k-mer modules

The pangenome and *k*-mer modules in PathoGD employ distinct methods to identify potential gRNA sequences within the target genome space. In the pangenome module, highly conserved protein-coding genes (≥90% prevalence across target genomes) are identified as potential targets for gRNA and primer design. Following a sequence clustering step to exclude non-target-specific genes *(see Methods)*, a consensus sequence alignment is obtained for each gene, from which canonical TTTN PAM sites are identified. The sequence downstream of the PAM site of a user-specified length *k* is stored as a potential gRNA sequence, or the gene is discarded if no PAM site is present. In genes where multiple PAM sites exist, ten gRNAs per PAM site are randomly selected, resulting in a maximum of 40 gRNAs per gene. A subsequent sequence comparison step eliminates gRNAs that are present in any non-target genome with up to 2 mismatches.

While the pangenome approach identifies universal genes, it excludes noncoding regions of the genome, including intergenic regions, ribosomal RNA and other noncoding RNA. These non-translated regions make up an average of 12% of the bacterial genome (33, 34), comprise many regulatory elements with key functions (35) and potentially contain species-specific sequence signatures (36) that could be exploited for diagnostics. To this end, we developed the *k*-mer module, a gene-agnostic approach interrogating both coding and non-coding regions across the entire genome. In the *k*-mer module, all sequences of a user-specified length *k* in both target and non-target species, including the human genome, are enumerated. Sequences common between the target and non-target genomes are removed, allowing for up to 2 mismatches in the non-target genomes (*see Methods*). A second *k-*mer enumeration step is performed to retain only sequences that are unique to the target species, downstream of a PAM site, and present above a genome prevalence threshold as potential gRNAs.

In both modules, gRNAs with the potential to form hairpin structures are eliminated, and primers designed from the flanking upstream and downstream sequences for the remaining gRNAs. The primer design step is optional in the *k*-mer module, and can be excluded where pre-amplification is not required for the assay. An *in silico* PCR is subsequently performed against all target and non-target genomes to estimate the sensitivity and specificity of each primer set. The output of PathoGD is a tab-delimited file providing up to five matching RPA primer pairs for each gRNA, along with associated information including GC content, expected amplicon size, and potential cross-reactivity with off-target organisms. This comprehensive output provides users with the flexibility to apply further filtering based on criteria such as primer and gRNA prevalence across target genomes, average copy number, and potential off-target hits.

### Minimizing potential off-target activity

Off-target activity in CRISPR-based diagnostics can occur through a combination of non-specific activity at the RPA and CRISPR-Cas stages. Non-specific amplification can occur as RPA has been shown to tolerate up to 9 primer-template mismatches (37–39). Combined with the potential imperfect binding of gRNA to non-target amplicons (40) or background DNA, this may result in the activation of trans-cleavage and generation of off-target signal, thereby reducing assay specificity.

PathoGD aims to minimize the possibility of false positives at the sequence level by implementing multiple steps to identify potential primer or gRNA binding sites in non-target genomes. In the pangenome module, core genes that are highly similar between target and non-target species are excluded through the clustering of target and non-target protein sequences at a 90% average amino acid identity (AAI) threshold. Cas12a has been demonstrated to tolerate mutations in the PAM and partial mismatches between gRNA and target region which permit target binding and cleavage to varying degrees (30, 40). To account for possible trans-cleavage activity resulting from imperfect target binding, candidate gRNAs with less than 3 mismatches to any sequence in the non-target or human genome databases are eliminated through a *k-*mer comparison step *(see Methods)*. These steps refine the pool of gRNAs to those with minimal off-target effects before proceeding to primer design, reserving computational resources for only promising and specific candidates. Additionally, mismatches between the gRNA and target DNA that occur in the seed region-the first 6-7 bases downstream of the PAM site, are more deleterious to Cas12a (41, 42) compared to mismatches at PAM-distal positions. To distinguish between these two categories, PathoGD incorporates a separate field denoting gRNAs where mismatches to non-target DNA occur exclusively outside of the seed region to allow for the selection of more robust gRNAs where available.

The selection of a primer and gRNA combination often involves a compromise between sensitivity and specificity. To facilitate decision-making, additional information is provided in the output: i) the proportion of genomes estimated to amplify from each primer set across the target and non-target genome databases, allowing up to two and eight overall mismatches for the target and non-target genomes, respectively, ii) prevalence of gRNAs with no mismatches in the target genome database, iii) gRNAs whose mismatches occurred exclusively at PAM-distal positions where they are mostly tolerated (*see Methods*). We suggest the following criteria be used where possible for identifying a highly reliable set of primer and gRNA: ≥90% prevalence of primers and gRNAs in target genome database, exclusion of primer sets with up to eight mismatches to off-target genomes, exclusion of gRNAs where mismatches to non-target genomes occur exclusively at PAM-distal positions, and prioritization of TTTA, TTTC, and TTTG PAM sites, due to demonstrated lower activity of wild-type LbCas12a nuclease on TTTT PAM sites (43, 44), which we also confirmed **(Figure S2)**. We note that the above are intended as general guidelines and should be adapted for different organisms or specific experimental contexts where necessary.

### Computational performance of PathoGD for primer and gRNA design for five clinically relevant human pathogens

We used PathoGD to design primer and gRNAs for five pathogens of global significance, which collectively represent four different phyla and include both Gram-negative and Gram-positive bacteria: *N. gonorrhoeae, Treponema pallidum, Chlamydia trachomatis, S. aureus* and *S. pyogenes.* These pathogens are high priority global threats according to the World Health Organization (WHO) with urgent benchmarks in place for novel diagnostics (45), urgent vaccine development (46) and new antibiotics (47) for pathogen elimination. Primers and gRNAs were designed for each species using genomes obtained from the NCBI Assembly database (**Table S2)**, subsampled to 100 genomes each except for *T. pallidum* where 92 genomes were available. The designs were subsequently validated against a larger genome database for each species *(see Methods)*, containing sequences with global representation and spanning a period of nearly two decades.

To improve specificity for primer and gRNA design, we included both ‘target’ and ‘non-target’ genomic data from species belonging to the same genus as the target species **(Table S2)**, and human genomic data *(see Methods)*. All designed primers and gRNAs were subsequently validated against the complete, non-subsampled ‘target’ and ‘non-target’ database for each species. Each run was performed on a computational cluster comprising Intel(R) Xeon(R) Gold 6342 CPU @ 2.80GHz (Icelake) processors using 32 CPU and 64GB RAM. Excluding the genome download step, run times ranged from 44 min to 5 h 26 min for the pangenome module and 41 min to 3 h 15 min for the *k*-mer module, and were largely correlated with the size of the target and non-target genome databases **(Table S3)**. Pipeline bottlenecks occurred at four stages: alignment of target genes or amplicons; enumeration of *k*-mers for non-target genomes; evaluation of guide specificity, and target gene annotation (specific to the pangenome module). Across the five species, these three steps accounted for 74.2 ± 5% and 79.1 ± 15% (mean ± standard deviation (SD)) of the total run times for the pangenome and *k*-mer modules, respectively. The computational runtime of PathoGD was significantly improved by the incorporation of tools employing efficient algorithms, including GenomeTester4 (48) for enumerating *k*-mers and manipulating the resulting lists, Primer3 (49) for primer design and evaluation, and isPcr (50) for *in silico* PCR amplification.

### Identification of primers and gRNAs using PathoGD

The number of target-specific gRNAs identified by PathoGD varied across the five species **(Figures 1a and 1c).** Despite having the largest genome sizes among the five species, *S. aureus* (∼2.83 Mbp) and *N. gonorrhoeae* (∼2.15 Mbp) had the least number of identified gRNAs across both pangenome and *k*-mer modules – 1,017 for *S. aureus* and 409 for *N. gonorrhoeae* **(Figure 1a,c; Figure S3; Table S3)**. In contrast, the two species with the smallest average genome sizes (∼1.1 Mbp) had the highest numbers of gRNAs – 10,035 for *C. trachomatis* and 15,157 for *T. pallidum,* respectively **(Figure 1a,c; Figure S3; Table S3)**, consistent with their relatively lower genetic diversity and highly adapted lifestyles (51–53). Overall, 5,317 gRNAs were identified for *S. pyogenes* from a smaller number of core genes compared to *S. aureus* and *N. gonorrhoeae*. These results translate to a higher number of actual primer/gRNA combinations available, given that up to 5 primer pairs were reported for each gRNA. Additionally, the above results were derived from observations where gRNAs and their associated primer pairs exhibited perfect matches to ≥90% of genomes in the complete target database, underscoring the utility of PathoGD in high-throughput identification of potential assay targets.

**Figure 1.**
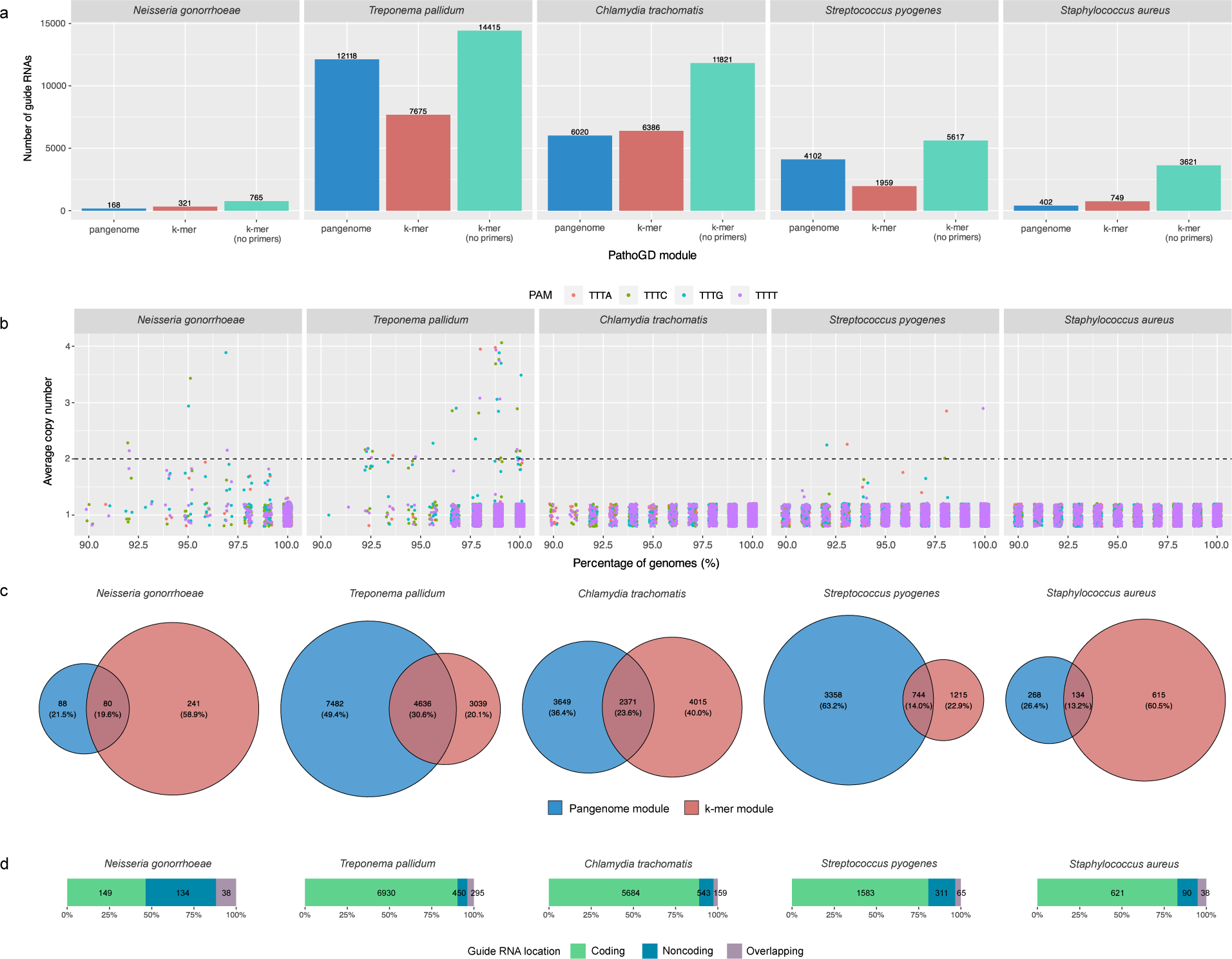
PathoGD *in silico* results of pangenome and *k*-mer modules for five bacterial species. **a)** Number of target-specific guide RNAs identified from the pangenome and *k*-mer modules in PathoGD. The *k-*mer (no primers) module refers to the workflow without primer design (*see Figure S1*). Results for the pangenome and *k*-mer modules were filtered to only observations where gRNAs and primers were present across 90% of strains in each target species, primers had at least nine mismatches to the non-target species, and at least one mismatch of gRNA to non-target genomes occur in the PAM or seed region. **b)** Average copy numbers of gRNAs identified from the *k*-mer (no primer) module for the five bacterial species. Black dotted lines refer to the cut-off at 2 copies. **c)** Unique and overlapping gRNAs identified from the pangenome and *k*-mer modules in (a) for five bacterial species. **d)** Distribution of gRNAs across the coding, noncoding or overlapping regions within the genome. All results were generated using the “user_all_subsample” workflow, with 100 genomes randomly subsampled for primer and guide RNA design for each target species, except for *T. pallidum* (*n*=92). The subsampled genomes used in both modules are identical for each target species.

Only partial overlaps of gRNAs were observed between the pangenome and *k*-mer modules (**Figure 1c**) due to the different algorithms employed by each module at the initial candidate gRNA identification steps *(see Methods)*. Specifically, in the pangenome module, heuristics employed for narrowing down the initial search space, including core gene elimination via average AAI to non-target genes and subsampling of a maximum of 10 gRNAs per canonical PAM for each gene, mean that certain gRNA candidates will be missed. In the *k*-mer module, *k*-mers are shortlisted only if present above a certain genome prevalence threshold, and if their flanking regions are sufficiently conserved. For the *k*-mer module, the percentage of gRNAs belonging to or overlapping with noncoding regions were estimated to range from 9.7 to 53.6% for the five species tested **(Figure 1d)**. These results, along with the substantial proportions of gRNAs unique to each module, support the complementarity of the pangenome and *k*-mer approaches in designing CRISPR-Cas12a diagnostic assays.

A further distinction between pangenome and *k*-mer modules in PathoGD is that the latter allows for the identification of target-specific sequences that occur in multiple copies throughout the genome. These sequences are attractive candidates for diagnostic targets in samples where low pathogen abundance is an issue, due to their potential for enhancing assay sensitivity. Additionally, the risk of false negatives is minimized in cases where certain copies are absent or have undergone mutations. The use of multi-copy markers as targets in molecular assays is not uncommon, for example IS2404 in *Mycobacterium ulcerans* (54), IS481 in *Bordetella pertussis* (55) and IS711 in *Brucella* spp. (56). As part of the *k-*mer module, multi-copy target-specific sequences are identified in an intermediate step of the pipeline and is the final output for a workflow not requiring primer design **(Figure S1)**. We identified 32 gRNAs with an average of ≥2 copies across *T. pallidum* (24 gRNAs), *N. gonorrhoeae* (4 gRNAs) and *S. pyogenes* (4 gRNAs) in our dataset **(Figure 1b)**; however, conserved primer binding sites were identified for only 14 of these (all in *T. pallidum*) likely due to polymorphisms in the gRNA flanking regions. The actual copy numbers of these individual gRNAs are highly variable within strains of a species **(Figure S4)** and results will likely differ when using a different subset of genomes.

### Comparison of PathoGD to PrimedSherlock

Currently, there is no reported pipeline for designing both primers and gRNAs for CRISPR-Cas12a diagnostic applications. However, a previously published tool, PrimedSherlock (29), described the design of gRNAs using primers generated from PrimedRPA (31), although published data are, to date, only available for viral genomes. Using the input target and off-target genome sequences reported in PrimedSherlock, we designed primers and 20-nucleotide gRNAs for 3 species of *Flavivirus*-West Nile virus (WNV), Zika virus (ZIKV), and four serotypes of Dengue virus (DENV-1, DENV-2, DENV-3, and DENV-4) using the PathoGD *k*-mer module, without genome subsampling. We compared the results of PathoGD to that of PrimedRPA and PrimedSherlock in terms of the potential to amplify their genomic regions and the prevalence and conservation of primer and gRNA binding sites across target and off-target genomes calculated using the same approach ***(see Methods)***.

The small average genome sizes (∼11kb) of these flaviviruses translated to few gRNAs, with less than 15 gRNAs obtained for DENV-1, DENV-2, DENV-4 and ZIKV, and 21 and 83 gRNAs for DENV-3 and WNV, respectively **(Figure S5a)**. Across all viruses, gRNAs from PathoGD had comparable or higher prevalence and conservation across the target genomes compared to PrimedSherlock, both when considering perfect matches and when allowing at most one mismatch between the gRNA and target genomes **(Figure S5b)**. Unsurprisingly, the prevalence of designed gRNAs was lower in these viruses compared to bacterial genomes, due to their high genetic diversity **(Figure S5b).** Only two gRNAs were common between PathoGD and PrimedSherlock-one for DENV-3 and one for WNV, present across ∼92% genomes in their respective databases. All gRNAs were specific to their target viruses, with only one DENV-1 gRNA from PrimedSherlock matching DENV-3 genomes at all but one position at the 3’ end. The primer binding sites were less conserved for dengue viruses compared to ZIKV and WNV, with no PathoGD primer candidates present in more than 50% of target genomes for DENV-1 and DENV-2 at a strict threshold of no allowed mismatches **(Figure S6a)**. However, allowing for mismatches across the forward and reverse primers increased the number of available primer candidates **(Figure S6a)** without compromising *in silico* primer specificity **(Figure S6b)**. Although the DENV-3 and DENV-4 primers reported in PrimedSherlock were more highly conserved **(Figure S6a; Figure S7a)**, PathoGD candidates had better overall prevalence and conservation for all six viruses when considering the primers and gRNAs in combination **(Figure S7)**.

### Experimental validation of primers and guide RNAs

To experimentally validate the primers and gRNAs designed *in silico*, we performed RPA-CRISPR-Cas12a assays for *S. pyogenes* and *N. gonorrhoeae*, following a previously established protocol (13). Seven sets of primer/gRNA candidates from *S. pyogenes* and six from *N. gonorrhoeae* present in ≥90% of genomes from their respective databases were selected from the list of primer and gRNAs generated from the PathoGD pangenome module **(Figure 2a**; **Figure 3a; Table S4)**. Additionally, primers and guides were manually designed for the widely used *N. gonorrhoeae* diagnostic target *porA* **(Table S4)**.

**Figure 2.**
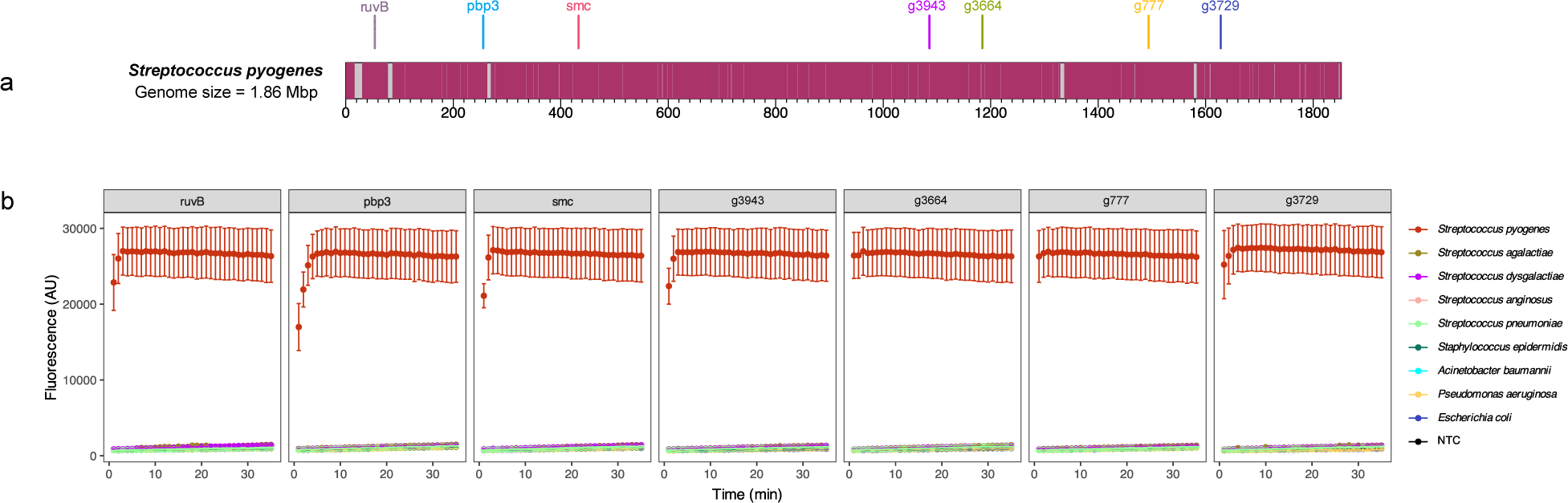
Experimental validation of primers and gRNAs for *Streptococcus pyogenes* using a previously developed RPA/CRISPR-Cas12a assay. **a)** Distribution of guide RNAs across the *S. pyogenes* ATCC 19615 genome (RefSeq assembly accession GCF_000743015.1). **b)** Fluorescence profiles of guide RNAs across 35 minutes for the *S. pyogenes* ATCC 19615 strain. Data represent average fluorescence values of three experimental replicates. Error bars represent SEM. The specificity of each gRNA was tested against a panel of closely related organisms and other organisms likely to be present at the site of infection.

**Figure 3.**
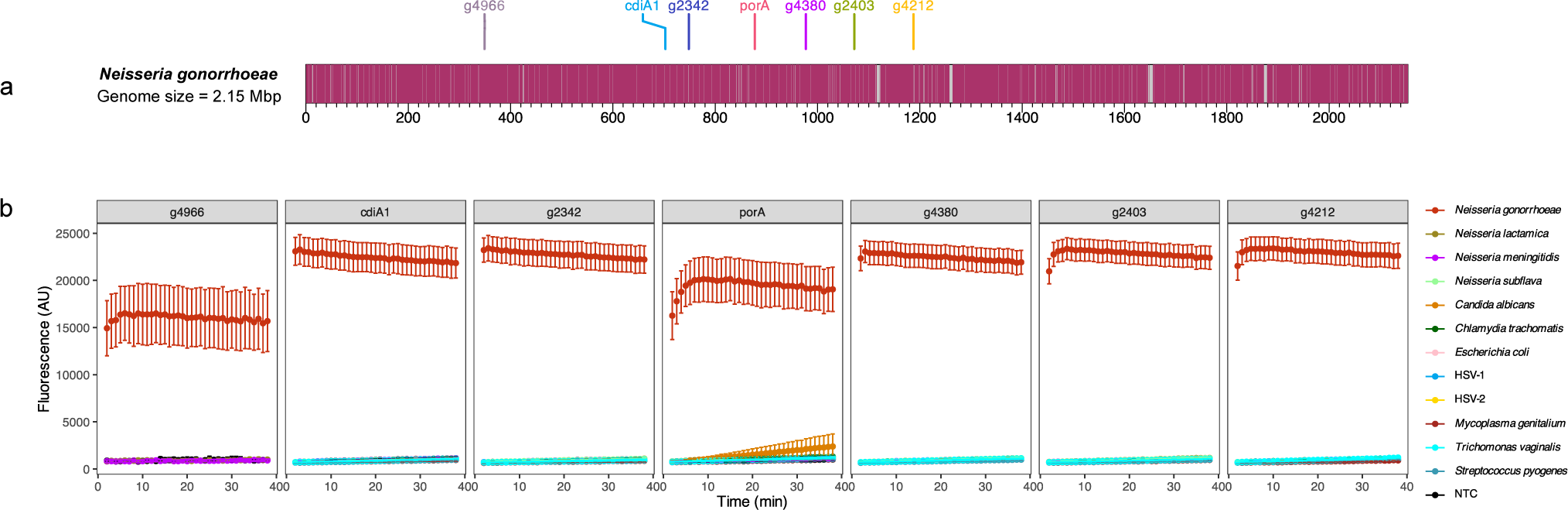
Experimental validation of primers and gRNAs for *Neisseria gonorrhoeae* using a previously developed RPA/CRISPR-Cas12a assay. **a)** Distribution of guide RNAs across the *N. gonorrhoeae* FA1090 genome (RefSeq assembly accession GCF_000006845.1) **b)** Fluorescence profiles of guide RNAs across 38 minutes for *N. gonorrhoeae* FA1090 strain. Data represent average fluorescence values of three experimental replicates. Error bars represent standard error of mean (SEM). The specificity of each gRNA was tested against a panel of closely related organisms and other organisms likely to be present at the site of infection.

All seven primer/gRNA sets detected *S. pyogenes* at 10,000 copies/μL and exhibited no cross-reactivity with genomic DNA of other species belonging to the same genus. These include the closely related *S. dysgalactiae* and *S. agalactiae*, as well as *S. anginosus* and *S. pneumoniae* **(Table S4; Figure 2b)**. In all instances, the fluorescence signals plateaued within the first five minutes, suggesting maximal reporter cleavage efficiency by Cas12a for the starting DNA concentrations used. A similar result was obtained for *N. gonorrhoeae*, where all six primer/gRNA sets accurately detected *N. gonorrhoeae* at 10,000 copies/μL. Only minimal background fluorescent signal was observed for a panel of closely related species, including *N. meningitidis* and *N. lactamica* (**Figure 3b**). In one of three runs, a minor fluorescence signal was observed in *Candida albicans* for the *porA* gRNA 20 minutes into the CRISPR/Cas12a assay, suggesting a possible contamination event (**Figure 3b**). Together, these results highlight the robustness of the primers and gRNAs designed by PathoGD.

In addition, we evaluated the potential of using multi-copy gRNAs for amplification-free detection of *N. gonorrhoeae*. Our amplification-free assay was based on an input gDNA of 250ng/µL, determined from previous experimental evaluation (data not shown) to be the minimum amount needed for detectable signal above background levels. From an initial assessment of 6 multi-copy gRNAs (average copy number ≥1.5) present in ≥90% of the target genomes **(Figure 4a; Table S4)**, four non-overlapping candidates with the highest fluorescent signals **(Figure 4b)** were further screened in an amplification-free CRISPR-Cas12a assay. These gRNAs were assessed individually, and incrementally pooled to generate 11 possible pooled combinations **(Figure 4c)**. These four gRNAs are present in 3 to 9 copies within the FA1090 genome, and cumulatively target 22 regions when pooled in a single reaction **(Figure 4a-b)**.

**Figure 4.**
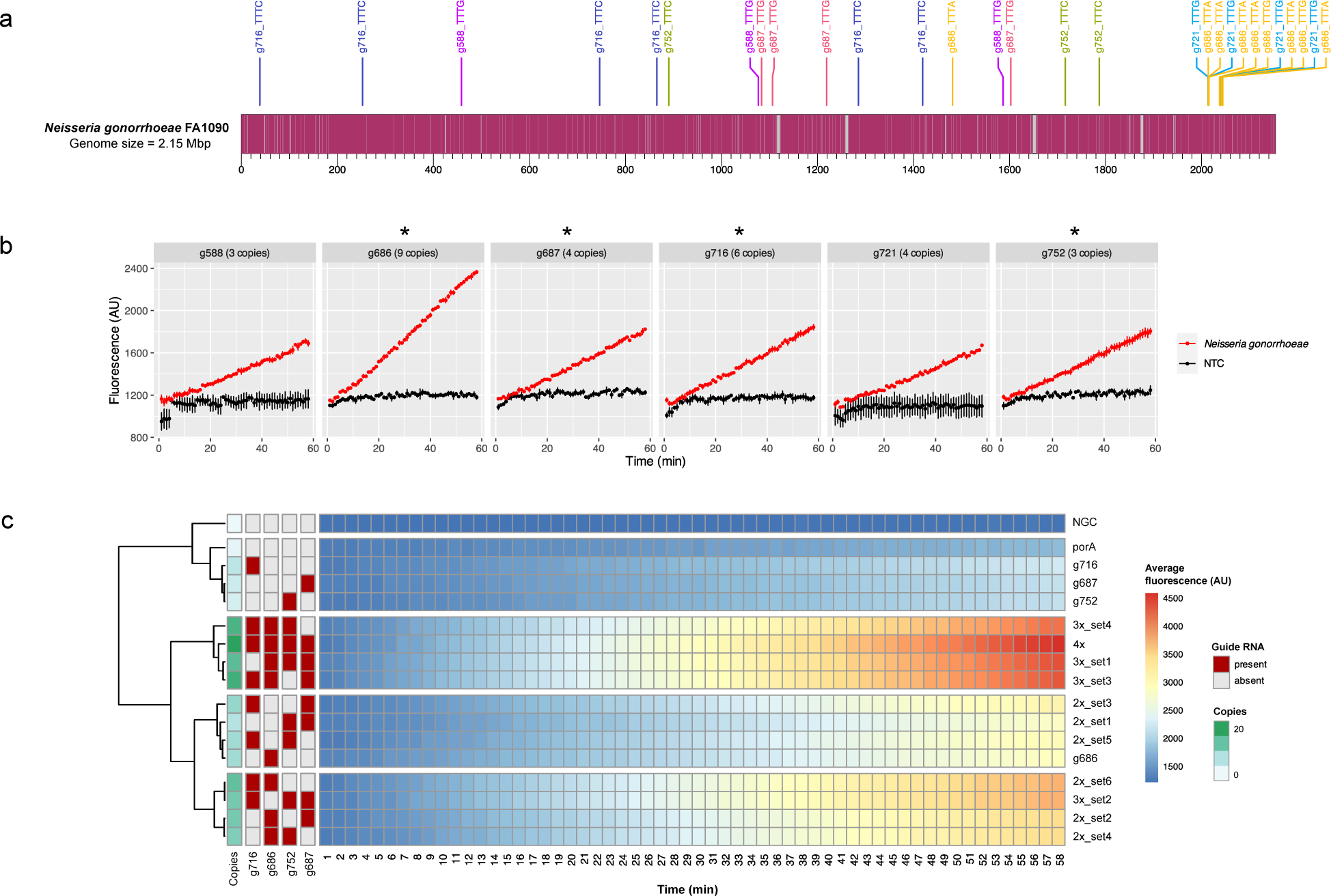
Experimental validation of gRNAs designed using *k-*mer module for *Neisseria gonorrhoeae*. **a)** Distribution of six selected multi-copy gRNAs across the *N. gonorrhoeae* FA1090 genome. Guide RNAs are present in multiple copies throughout the genome and denoted by their names and PAM sequence separated by an underscore. **b)** Initial assessment of six multi-copy gRNAs on *N. gonorrhoeae* FA1090 gDNA in an amplification-free CRISPR-Cas12a assay. The copy number for each gRNA in the FA1090 genome is listed next to the gRNA names. Data represent average fluorescence signal across 58 min. Error bars represent SEM across three experimental repeats (two technical replicates for *N. gonorrhoeae* in each experimental repeat). The four gRNAs selected for pooling are indicated with an asterisk. **c)** Heatmap of average fluorescence profiles for individual and pooled multi-copy gRNAs in an amplification-free CRISPR-Cas12a assay using four selected gRNAs. NGC refers to the control with no guide RNA added. Fluorescence values were averaged across multiple experimental repeats (*n* = 3 for porA, *n* = 6 for all other gRNA sets). The presence-absence profiles of individual gRNAs across all individual and pooled combinations tested and their cumulative copy numbers in the FA1090 genome are annotated on the left of the heatmap.

All multi-copy gRNAs successfully detected *N. gonorrhoeae*, albeit with a much slower signal accumulation compared to an assay with RPA preamplification. As expected, combinations of gRNAs with higher cumulative copy numbers resulted in higher endpoint fluorescence, with the signal in most combinations being driven by g686, the best-performing gRNA **(Figure 4b-c)**. Pooled multi-copy gRNAs resulted in at least 1.5-fold increase in fluorescence signals at the assay endpoint of 58 min compared to *porA*, a single copy marker **(Figure S8-9)**. The fluorescence signals did not plateau for any of the gRNA sets **(Figure S8)**, even when the assay was extended to 118 minutes **(Figure S8)**, consistent with previous observations (57) and suggesting suboptimal activation of Cas12a in a preamplification-free assay. Encouragingly, however, multi-copy gRNAs were highly specific, with only minimal off-target signal observed in the presence of 4 pooled gRNAs beginning at 20 minutes **(Figure S8-9)**.

## DISCUSSION

Critical to the success of CRISPR-based diagnostics is the design of highly sensitive and specific primers and gRNAs for target detection, a process generally involving multiple iterative steps. Here, we describe PathoGD, an end-to-end bioinformatic pipeline for the high-throughput and rapid design of highly conserved and specific primers and gRNAs for CRISPR-Cas12a diagnostic applications. PathoGD integrates multiple tools employing efficient algorithms and requires only a user-specified configuration file as input, significantly reducing the complexity and time required for assay design. PathoGD is reliant on a well-curated target (inclusivity) and non-target (exclusivity) genome database to capture the full spectrum of genetic diversity within the target species while excluding closely related species and organisms likely to co-occur at the site of infection. We demonstrated the utility of PathoGD for designing primers and gRNAs for five diverse species of bacterial pathogens. Computational runtimes were largely dependent on database sizes, with the longest runtime observed for *S. pyogenes* (5h 26 min) for the pangenome module; however, all other runs were completed within 4 hours for both modules. Overall, the use of a small subset of 100 genomes per species was sufficient to generate an extensive set of potential targets that were valid across the larger complete target database.

PathoGD eliminates the reliance on traditionally used marker genes when selecting a target region for CRISPR-based detection by leveraging publicly available genome assemblies. In addition to a gene-centric (pangenome) approach, the incorporation of a complementary gene-agnostic (*k*-mer) approach allows for a systematic and unbiased exploration of sequences in both protein-coding and non-translated regions of the genome for more challenging organisms. As the field of molecular diagnostics is transitioning towards a dual-target approach to minimize the risk of false negatives, an essential step is the evaluation of multiple candidate primers and gRNAs as their sensitivities and specificities *in vitro* may vary. The comprehensive output generated by PathoGD provides flexibility in the selection of candidates for experimental validation. Equally important is the continuous monitoring of newly sequenced genomes of pathogens for potential mutations that could result in diagnostic escape. The scalability of PathoGD ensures that the design process can easily be revisited with the newly sequenced genomes included, or alternatively, previously designed primers and gRNAs can be evaluated against new genomes to validate their ongoing relevance.

The number of target-specific gRNAs identified by PathoGD varied across species and could be attributed to the following factors: core genome size, allelic variants within core genes, evolutionary relationship with closely related species, and the genetic nature and diversity of strains subsampled for each species. Organisms sharing large homologous regions with closely related species and frequently engaging in horizontal gene transfer present a challenge for target-specific primer and gRNA design. In our dataset of five species, this was exemplified by *N. gonorrhoeae* which shares between 80 and 90% of its genome sequence with its closest relative *N. meningitidis* (58), is naturally competent for DNA uptake (59, 60), undergoes frequent recombination with other *Neisseria* species (61), and exhibits significant genetic variation between subtypes (62). These factors contribute to complexity in marker selection for accurate and specific detection of *N. gonorrhoeae*, especially for specimens from extragenital sites where commensal *Neisseria* species are commonly found (62). Indeed, commercial gonococcal nucleic acid amplification tests (NAAT) have displayed variable cross-reactivity with other *Neisseria* species and often require a supplementary assay for confirmation (63). Across both pangenome and *k*-mer modules, PathoGD identified a total of 409 *N. gonorrhoeae*-specific gRNAs from 31 genes. Our experimental validation on a subset of these genes supports their potential as alternative targets for specific detection of *N. gonorrhoeae* that could be assessed for increased sensitivity at low pathogen concentrations, compared to *porA* which has been utilized by *N. gonorrhoeae* CRISPR-Cas assays to date (18, 64, 65).

Despite its smaller proportion of core genes compared to *N. gonorrhoeae*, we identified a substantially larger number of gRNAs (*n* = 5,317) in *S. pyogenes* due to a higher number of species-specific genes. Unlike *porA* in *N. gonorrhoeae*, there are limited data regarding a widely used NAAT diagnostic target for *S. pyogenes*. We validated a subset of seven primer and gRNA combinations for the detection of *S. pyogenes* and demonstrated high specificity, highlighting the reliability of primers and gRNAs designed by PathoGD. Although only a single strain of target and non-target species was used for all validation assays, *in silico* evaluation against a large database of closely related species suggested that the gRNAs are robust for discriminating target and non-target organisms with a broad range of genetic diversity.

Across other assessed species, varying numbers of gRNAs were identified as a product of their genetic diversity, evolutionary history, ecological niche and degree of relatedness to other species. Importantly, the targets we identified were validated across comprehensive databases with sequences that have wide geographical distribution and have persisted over evolutionary time scales, translating into increased longevity and robustness of the diagnostic assay. In our example runs, the off-target databases consisted only of species belonging to the same genus as the target pathogen, which may be overly conservative considering that many may not be present in the same sites or environment as the target pathogen. Likewise, the inclusion of more distantly related species in the off-target database should be carefully considered for organisms where horizontal gene exchange across genera or higher taxonomic ranks is prevalent.

The lack of overlap between gRNAs generated from PathoGD and PrimedSherlock can be attributed to their different underlying approaches. Specifically, PathoGD starts by identifying potential gRNAs which serve as anchors for primer design, while PrimedSherlock designs gRNAs from pre-defined genomic regions amplified by user-provided primer sequences. Additional considerations in PathoGD which were absent in PrimedSherlock include elimination of gRNA candidates with homopolymers of five or more nucleotides, as well as those matching the human genome. These highlight the currently evolving field of CRISPR diagnostics where no consensus exists yet for standard approaches in primer and gRNA design. Our results with the DENV-3 dataset suggested that some pipeline parameters may have to be relaxed when designing assays for viruses to increase the number of available primer and gRNA candidates, especially for rapidly mutating RNA viruses. The simultaneous design of primers and gRNAs is also advantageous for upfront elimination of genomic regions that do not contain PAM sites or are poorly conserved across the target species.

Previous studies have shown success in preamplification-free detection of SARS-CoV-2 (57) and hepatitis C virus (66) with Cas13a and *Mycobacterium ulcerans* (*66*) with Cas12a. The availability of candidate gRNAs present in multiple copies across the genome in *N. gonorrhoeae* presented an opportunity to evaluate the potential increase in detection sensitivity through the use of multi-copy gRNAs, both individually and pooled. Our experiment supported previous studies reporting an increased sensitivity with the inclusion of more gRNAs, with a clear relationship observed between signal accumulation and cumulative regions targeted by the gRNAs. While not sufficient to detect pathogens at low concentrations on its own, the pooling of highly specific multi-copy non-overlapping gRNAs may be valuable when used in conjunction with post signal amplification approaches compatible with CRISPR-Cas assays (67, 68). The presence of multi-copy species-specific gRNAs in bacterial genomes is not ubiquitous, as demonstrated by the *in silico* results for the five species, but can be exploited where available as the CRISPR diagnostics field shifts towards amplification-free methods for nucleic acid detection. In this regard, PathoGD can address the need for rapid identification of highly specific multi-copy potential diagnostic targets.

There are several limitations of PathoGD, some of which can be addressed in future versions of the tool. First, gRNA designs in PathoGD are limited to those adjacent to Cas12a-preferred TTTN PAM sites. Future improvements could expand its utility for other Cas nucleases with different PAM requirements. Second, the success of a run is dependent on the taxonomic accuracy of genomes in the target and non-target databases. A single contamination in either database due to taxonomic or assembly anomalies is sufficient to remove or significantly reduce the number of potential target-specific gRNAs identified. This is due to the *k*-mer comparison step employed by PathoGD, where sequences present in both databases would be eliminated. A future version would benefit from an integrated taxonomic validation step. Finally, PathoGD refrains from ranking or scoring the designed primers and gRNAs, as their binding and cleavage efficiencies are dependent on multiple factors that may not be fully captured by a simple scoring system. Studies focused on CRISPR editing applications in eukaryotes have shown that gRNA sequence composition, chromatin accessibility, and RNA secondary structure and free energy contribute to gRNA efficiency (69). Future large-scale studies investigating the above factors could pave the way for incorporating machine learning in predicting gRNAs with maximal activity for bacterial diagnostics, such as that implemented with Cas13 for viruses (70).

## CONCLUSIONS

The emergence of CRISPR-based pathogen diagnostics as promising alternative in molecular diagnostics has necessitated the development of bioinformatic pipelines capable of rapid assay design. PathoGD addresses the challenges associated with the design of highly specific primers and gRNAs for target organisms of interest using the wealth of publicly available genomic data. The integration of computationally efficient tools in PathoGD enables rapid selection of potential diagnostic targets, and the generation of a comprehensive list of primers and gRNAs provides numerous alternatives for laboratory evaluation and validation. We anticipate that PathoGD will contribute to the accurate and efficient design of CRISPR-Cas12a-based diagnostic assays as the field continues to expand. PathoGD is available open-source on GitHub at https://github.com/sjlow23/pathogd.

## METHODS

### Pangenome module

To identify a set of targets for amplification, gene annotation was performed using Prokka (v1.14.6) (71) for all target or subsampled target genomes, depending on user input. The pangenome was estimated using Roary (v3.13.0) (72) for the same set of target genomes. Conserved genes present in at least 90% of all genomes were identified as potential targets for amplification. Structural gene annotation was performed for all non-target genomes using Prodigal (v2.6.3) (73). Protein sequences of conserved target and non-target genomes were clustered at 90% amino acid identity threshold using cd-hit (v4.8.1) (74), and only gene families from target genomes which did not cluster with genes from non-target genomes were retained as target-specific genes. All target-specific gene families were individually aligned using MAFFT v7.505 (75), and a consensus gene alignment obtained using EMBOSS (v6.6.0; http://emboss.open-bio.org) as the template for gRNA and primer design. Canonical PAM sites (TTTA, TTTC, TTTG, TTTT) were identified from each consensus gene alignment on both sense and antisense strands. Sequences downstream of the PAM site of a user-specified length *k* were stored as potential gRNA sequences, or the gene was discarded if no PAM sites were present. Up to 10 potential gRNA sequences were retained per PAM site to give a maximum (31) of 40 gRNA sequences per gene. Low-quality potential gRNAs were removed based on their propensity to form hairpin structures using the ‘check_primers’ task from Primer3 (v2.6.1) (49). A *k*- mer comparison was performed between the remaining potential gRNA sequences and *k*-mers of the same length enumerated from non-target genomes using GenomeTester4 (v4.0) (48) to eliminate gRNAs present in the non-target genomes, allowing at least 2 mismatches. Flanking sequences of up to 150 bp (where available) were extracted for the remaining gRNA sequences using SeqKit (v2.3.1) (76), and primers designed for an amplicon size range of 90 – 300 bp using Primer3. A maximum of 5 primer pairs were designed for each gRNA. An *in silico* PCR was performed against all target and non-target genomes with the designed primers using isPcr (v33) (50) to estimate the prevalence of each target amplicon across the target and non-target genome databases. Similarly, all gRNAs were mapped to target and non-target genomes using BBMap (v38.96) (77) to obtain an estimate of gRNA prevalence.

### k-mer module

*k*-mers of a user-specified length were enumerated for all target or subsampled target genomes and all non-target genomes using GenomeTester4, followed by a *k*-mer comparison between the target and non-target genomes to eliminate potential gRNA sequences with up to 2 mismatches in the non-target genomes. A second *k*-mer enumeration step of length *k*+4 was performed for the target genomes to identify target-specific *k*-mers with PAM sites. Low-quality potential gRNAs from this list were removed based on their propensity to form hairpin structures using the ‘check_primers’ task in Primer3. The target-specific *k*-mers were mapped to the target genomes using BBMap and flanking sequences of up to 150 bp (where available) were extracted for each *k*-mer as potential amplicons. All amplicons were clustered using cd-hit, and clusters above 90% nucleotide identity were individually aligned. Primers were designed each consensus alignment for an amplicon size range of 90 – 300 bp. A maximum of 5 primer pairs were designed for each gRNA. An *in silico* PCR was performed against all target and non-target genomes with designed primers using isPcr to estimate the prevalence of each target amplicon across target and non-target genome databases. Similarly, all gRNAs were mapped to target and non-target genomes using BBMap to obtain an estimate of gRNA prevalence.

### Data visualization

For visualization of gRNAs on bacterial genomes, gRNA sequences were mapped to one representative genome from each species using BLAST (v2.13.0) (78) with the flags “-qcov_hsp_perc 100 -perc_identity 100 -outfmt 6 -task blastn-short”. Schematics of guide locations on the genomes were generated using the karyoploteR (v1.26.0) (79) package in R. All other graphs were generated using the ggplot2 (v3.4.2) (80) package in R. To determine genomic locations of gRNAs within the genome, one genome representative was selected for each species and annotated using Prokka to obtain coordinates of protein-coding regions. Sequences of gRNAs were mapped against their respective representative genomes using BLAST to obtain their genomic coordinates. Overlaps between the gRNAs with the genic and inter-genic regions within each genome were determined using bedtools (v2.30.0) (81).

### In silico primer and guide RNA design

For *in silico* testing of the PathoGD pipeline on five pathogens using both pangenome and *k*-mer modules, non-overlapping genome assemblies at the complete, chromosome, and scaffold levels were downloaded from NCBI RefSeq and GenBank (accessed June 27, 2023) for the target database for *T. pallidum* (*n* = 92), *C. trachomatis* (*n* = 171) and *S. pyogenes* (*n* = 765) based on taxonomic annotations obtained from the Genome Taxonomy Database (GTDB) r214. Genome assemblies at all levels were retained for *N. gonorrhoeae* (*n* = 978) due to high genetic diversity between subtypes. Genomes were limited to complete and chromosome levels for *S. aureus* (*n* = 1,096) due to overrepresentation in the database. For the non-target database, genome assemblies at all levels were retained for *N. gonorrhoeae* (*n* = 2,597), *T. pallidum* (*n* = 642), *C. trachomatis* (*n* = 268) and *S. aureus* (*n* = 4,600), except for one non-target species for *S. pyogenes* (*n* = 7,057), where 1,000 genomes were subsampled from the total for *S. pneumoniae, S. agalactiae* and *S. suis*. Genomes of all other species belonging to the same genus as the target species were used as the non-target database, except for *T. pallidum* and *C. trachomatis* where GTDB taxonomic discrepancies with NCBI were observed at the genus level. For *T. pallidum*, all genomes belonging to “g__Treponema”, “g__Treponema_B”, “g__Treponema_C”, “g__Treponema_D”, and “g__Treponema_F” in GTDB r214 were used for populating the non-target database. For *C. trachomatis*, all genomes belonging to “g__Chlamydia” and “g__Chlamydophila” in GTDB r214 were used for populating the non-target database. Additionally, the human reference genome (NCBI RefSeq assembly GCF_000001405.40) was included as part of the non-target database for all *in silico* designs. Primers and guide RNAs were designed using the PathoGD pangenome and *k*-mer modules with the ‘user_all_subsample’ workflow. For each species, a common subset of 100 target genomes were used for primer and gRNA design in both modules. All designed primers and gRNAs for each species were validated *in silico* against their respective full target databases. The metadata for all genome assemblies used was obtained from NCBI.

### Comparison to gRNA designs from PrimedSherlock

Genomic sequences of Dengue virus serotype 1 (DENV-1; *n* = 2,230), Dengue virus serotype 2 (DENV-2; *n* = 776), Dengue virus serotype 3 (DENV-3; *n* = 946), Dengue virus serotype 4 (DENV-4; *n* = 214), West Nile virus (WNV; *n* = 2,618) and Zika virus (ZIKV; *n* = 557), and sequences from their respective non-target databases (DENV-1; *n* = 5,555, DENV-2; *n* = 6,206, DENV-3; *n* = 4,014, DENV-4; *n* = 3,580, WNV; *n* = 6,957, ZIKV; *n* = 9,164) and gRNA sequences were obtained from PrimedSherlock. Primers and gRNAs were designed for the six viruses using the *k*-mer module of PathoGD without genome subsampling, with a specified amplicon size range of 90 to 500 base pairs, a gRNA prevalence threshold of 50%, and with the cd-hit amplicon sequence identity clustering threshold set to 80%. For DENV-3, the “--mismatch 1” flag was used for GenomeTester4 in the *k*-mer comparison steps as no output was obtained for the default setting. Primers and gRNAs from PathoGD and PrimedRPA/PrimedSherlock for each species were evaluated based on prevalence and conservation of their binding sites across all target and off-target genomes. Specifically, primers were assessed based on their potential to amplify their respective genomic regions using isPcr with the “-minPerfect=5” and “-maxSize=500” options, and prevalence across target genomes calculated for exact matches, ≤2 mismatches, ≤5 mismatches and ≤8 mismatches across the forward and reverse primers. Guide RNAs were evaluated based on presence of the gRNA binding sites along with an upstream TTTN PAM in their respective target and non-target genome databases using GenomeTester4. For target genomes, gRNA comparisons were made with no mismatches (“--mismatch 0” flag) or one mismatch allowed (“--mismatch 1” flag). For non-target genomes, comparisons were made allowing one (“--mismatch 1” flag), two (“--mismatch 2” flag), or three (“--mismatch 3” flag) mismatches.

### Experimental validation of primers and guide RNAs with RPA-CRISPR-Cas12a assay

Genomic DNA (gDNA) from the target organisms, *N. gonorrhoeae* and *S. pyogenes*, as well as organisms in the specificity panel were amplified using RPA with the TwistAmp Basic kit (TwistDx) according to the manufacturer’s instructions. The approximate starting amounts for each organism ranged from approximately 9,500 to 68,000 copies/µL **(Table S5)**. The RPA pellet was first resuspended with rehydration buffer supplied in the kit containing 0.48 μM of forward primer, 0.48 μM of reverse primer, and nuclease-free water to a total volume of 41 μL. The resuspended volume was split equally across four reactions, and 1.0 μL of template DNA and 1.25 μL of magnesium acetate (140 mM stock) was added. The reactions were incubated at 37°C for 30 min in a T100 Thermal Cycler (Bio-Rad).

LbCas12a trans-cleavage assays were performed similarly to a previously described assay (13). A total of 40 nM LbCas12a (Gene Universal) was preincubated with 50 nM gRNA and 2 μL RPA product in 1x Holmes Buffer 2 in a black 96-well assay plate for 10 min at 37 °C (Hybaid hybridization oven). The /56-FAM/TTTTTTTTT/3BHQ-1/ reporter molecule (IDT) was added to the reaction at a final concentration of 125 nM. Fluorescence was measured using the CLARIOstar Plus Microplate Reader (BMG LABTECH) every minute for 38 min (λ_ex_ = 490/8 nm; λ_em_ = 520/8 nm) at 37°C. No-template controls (NTC) were included as negative controls in all assays and contained all reagents except for template DNA. All RPA and CRISPR-Cas12a assays were performed with three experimental repeats.

For the amplification-free assays, LbCas12a trans-cleavage reactions were performed as above, with the exception that 250 ng/µL of *N. gonorrhoeae* gDNA was added. For the initial assessment of 6 multi-copy gRNAs, the assay was run for 58 min with 3 experimental repeats. For the pooled gRNA evaluation, the assay was run for either 58 (*n*=5) or 118 min (*n*=1), with 6 experimental repeats for all gRNAs and 3 experimental repeats for *porA*. Due to the high starting concentration required, specificity testing was performed only for the *porA* gRNA and reactions containing 4 pooled gRNAs, using 250 ng/µL of *N. lactamica*, *N. meningitidis*, and *N. subflava*, *C. trachomatis*, *E. coli*, and *S. pyogenes* gDNA. *In silico* analysis of the gRNA sequences confirmed no hits were observed against the remaining organisms in the *N. gonorrhoeae* specificity panel in Table S5.

### Statistical analyses

All statistical analyses were performed using R version 4.2.2.

## Availability of data and materials

The datasets used and/or analysed during the current study are available in the Supplementary Tables file. The genome assemblies used for the *in silico* design of primers and guide RNAs for the five pathogens are available in Table S2. Sequences and metadata for the primers and guide RNAs designed *in silico* are available from the corresponding author upon request. PathoGD is available open-source on GitHub (https://github.com/sjlow23/pathogd).

## Authors’ contributions

D.A.W., S.P. and S.J.L. designed the study. D.A.W. and S.P. provided supervision and support. S.J.L. developed the pipeline and performed bioinformatic analyses, with input from G.T., L.W. and S.P. M.O., W.J.K., and N. D. performed the experimental validations, with contributions from M.K., F.A, Y.N., and E.H. E.S. provided expertise in software development. S.J.L., S.P. and D.A.W wrote the initial draft of the manuscript. All authors read and approved the final manuscript.

