## Supplementary Figures for "PathoGD: an integrative genomics approach for CRISPR-based target design of rapid pathogen diagnostics"

### Low et al.

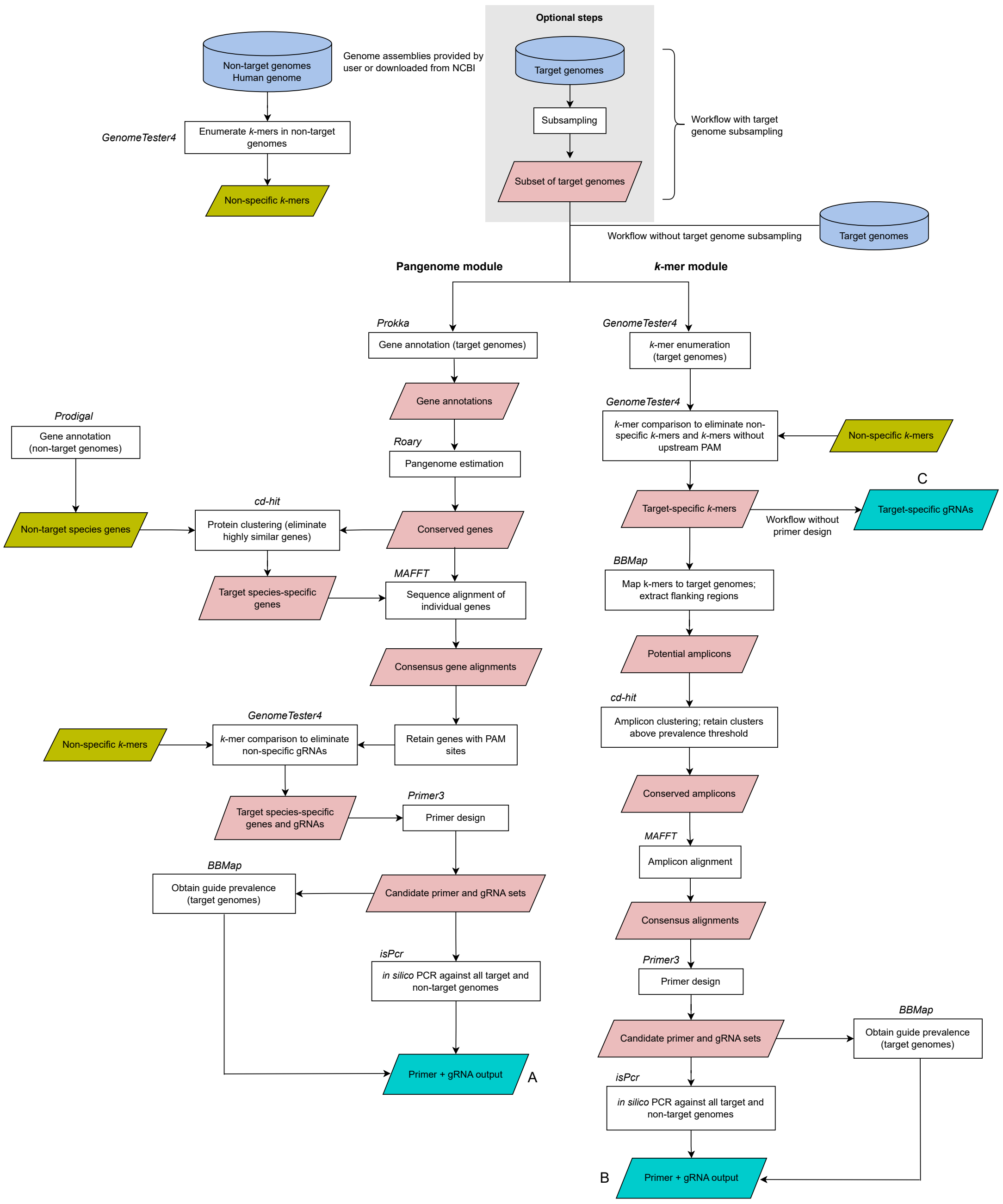

**Figure S1.** Overview of the main steps of the pangenome and *k*-mer modules in PathoGD. Users can provide their own custom set(s) of target and/or non-target genomes, or optionally automatically download from NCBI. The final output from the pangenome (A) and *k*-mer module (B) is a comprehensive list of target-specific primer and guide RNA combinations from which users can apply further filtering based on multiple criteria. The primer design step is optional in the *k*-mer module, without which a separate output of target-specific guide RNAs is generated (C).

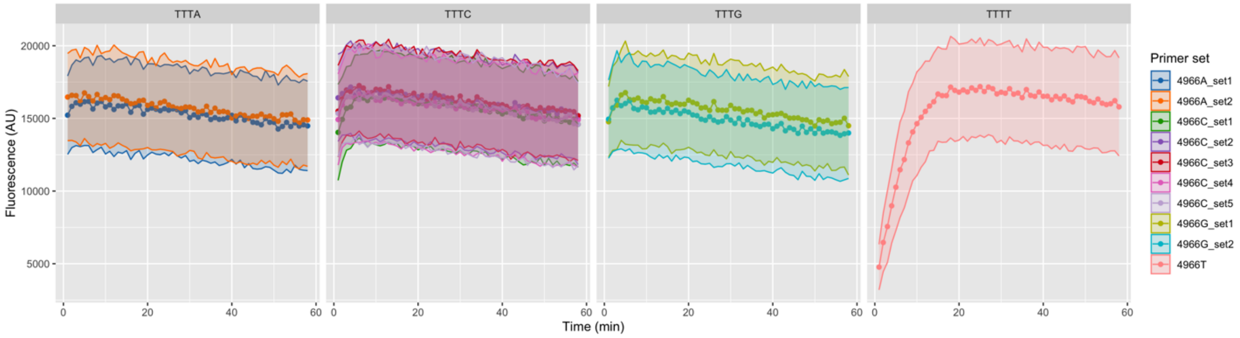


**Figure S2.** Fluorescence profiles for 4 gRNAs with TTTA, TTTC, TTTG and TTTT PAM sites designed for a single *N. gonorrhoeae* target gene used in conjunction with one or more RPA primer sets per gRNA. Data represent average fluorescence ± standard error of mean (SEM) across three experimental replicates. The starting concentration of *N. gonorrhoeae* DNA used was ~4.3 x $10^6$ genome copies/µL.

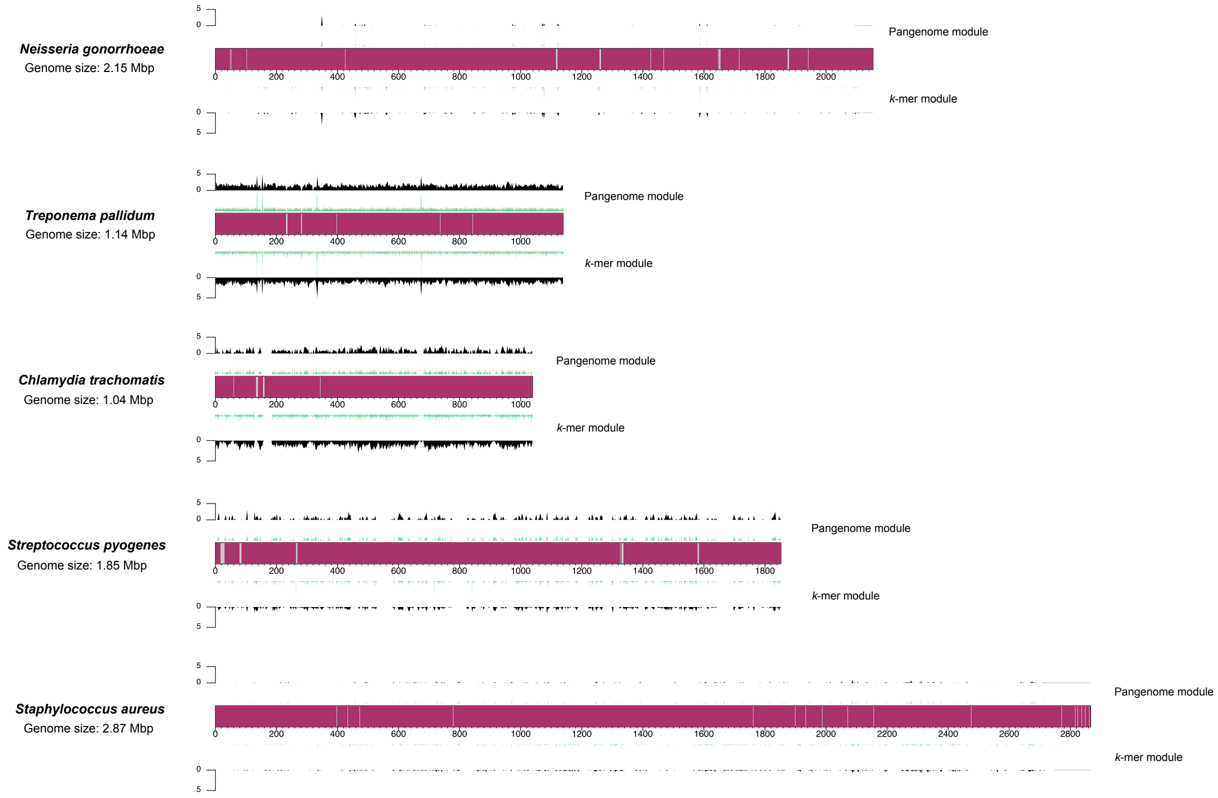


**Figure S3.** Distribution of guide RNAs designed by PathoGD across the genomes of five bacterial pathogens: *Neisseria gonorrhoeae* (NC_002946.2), *Treponema pallidum* (NC_010741.1), *Chlamydia trachomatis* (NC_021050.1), *Streptococcus pyogenes* (NC_002737.2) and *Staphylococcus aureus* (NZ_CP039167.1). Results for the pangenome and *k*-mer modules were filtered to only observations where gRNAs and primers were present across 90% of strains in each target species, primers had at least nine mismatches to the non-target species, and at least one mismatch of gRNA to non-target genomes occur in the PAM or seed region. Each bacterial chromosome is represented by a rectangle with a length proportional to its genome size; pink-shaded areas represent coding regions and gray-shaded areas represent non-coding regions. Each chromosome has a top and bottom panel representing gRNAs from the pangenome and *k*-mer modules, respectively. The first layer in each panel shows the mapped regions of the designed gRNAs and the black density plots on the second layer the number of gRNAs overlapping each 2,500 bp window.

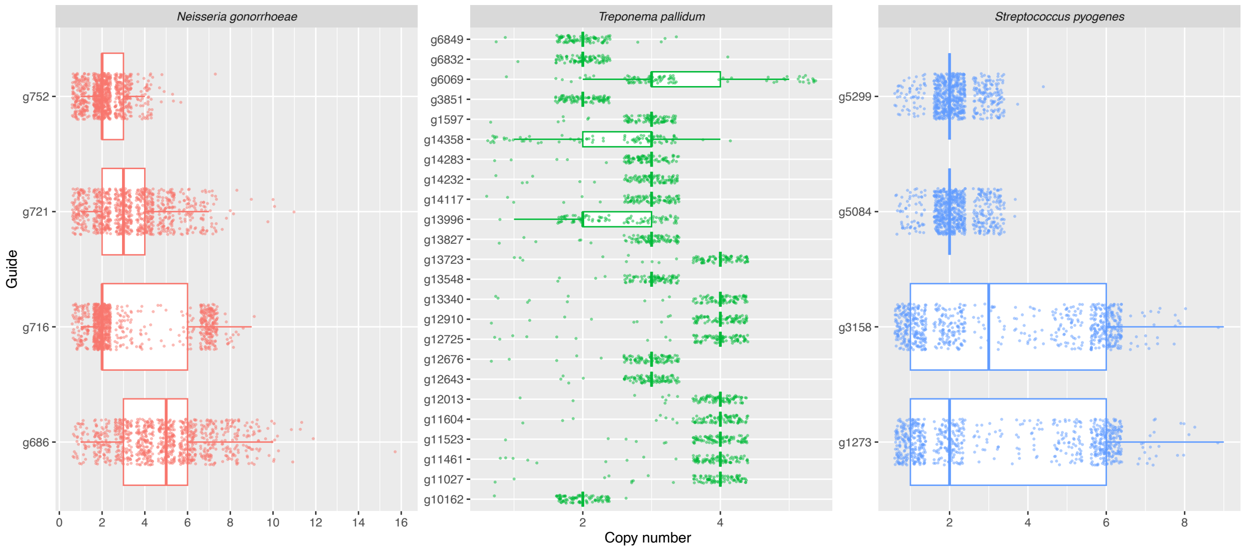


**Figure S4.** Distribution of copy numbers for a subset of guide RNAs across the individual genomes of *Neisseria gonorrhoeae* (*n* = 978), *Treponema pallidum* (*n* = 92), and *Streptococcus pyogenes* (*n* = 765). The subset of guide RNAs were designed by the PathoGD *k*-mer module and have an average copy number of ≥2 across the subsampled set of genomes for each species (*n* = 100 for *N. gonorrhoeae* and *S. pyogenes*; *n* = 92 for *T. pallidum*). The centre line of the boxplots corresponds to the median, the lower and upper hinges correspond to the first and third quartiles, respectively, and the ends of the whiskers represent values no further than 1.5x of the interquartile range.

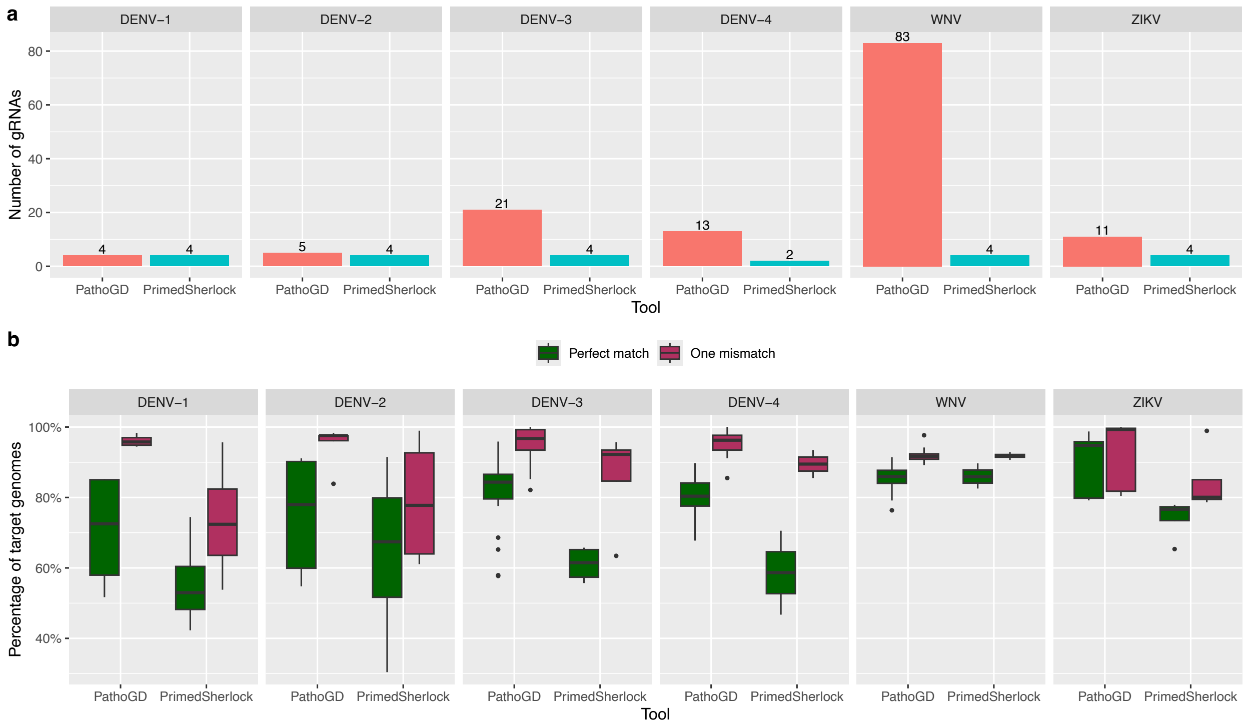


**Figure S5. Comparison of gRNAs designed by PathoGD and PrimedSherlock for six viruses. a)** Number of gRNAs designed using PathoGD and PrimedSherlock for six viruses. PrimedSherlock gRNAs were reported as the best candidates. **b)** Percentages of genomes targeted by gRNAs designed using PathoGD and PrimedSherlock for the six viruses, with perfect matches or at most one mismatch. DENV-1: Dengue virus serotype 1; DENV-2: Dengue virus serotype 2; DENV-3: Dengue virus serotype 3; DENV-4: Dengue virus serotype 4; WNV: West Nile virus; ZIKV: Zika virus. The centre line of the boxplots corresponds to the median, the lower and upper hinges correspond to the first and third quartiles, respectively, and the ends of the whiskers represent values no further than 1.5x of the interquartile range.

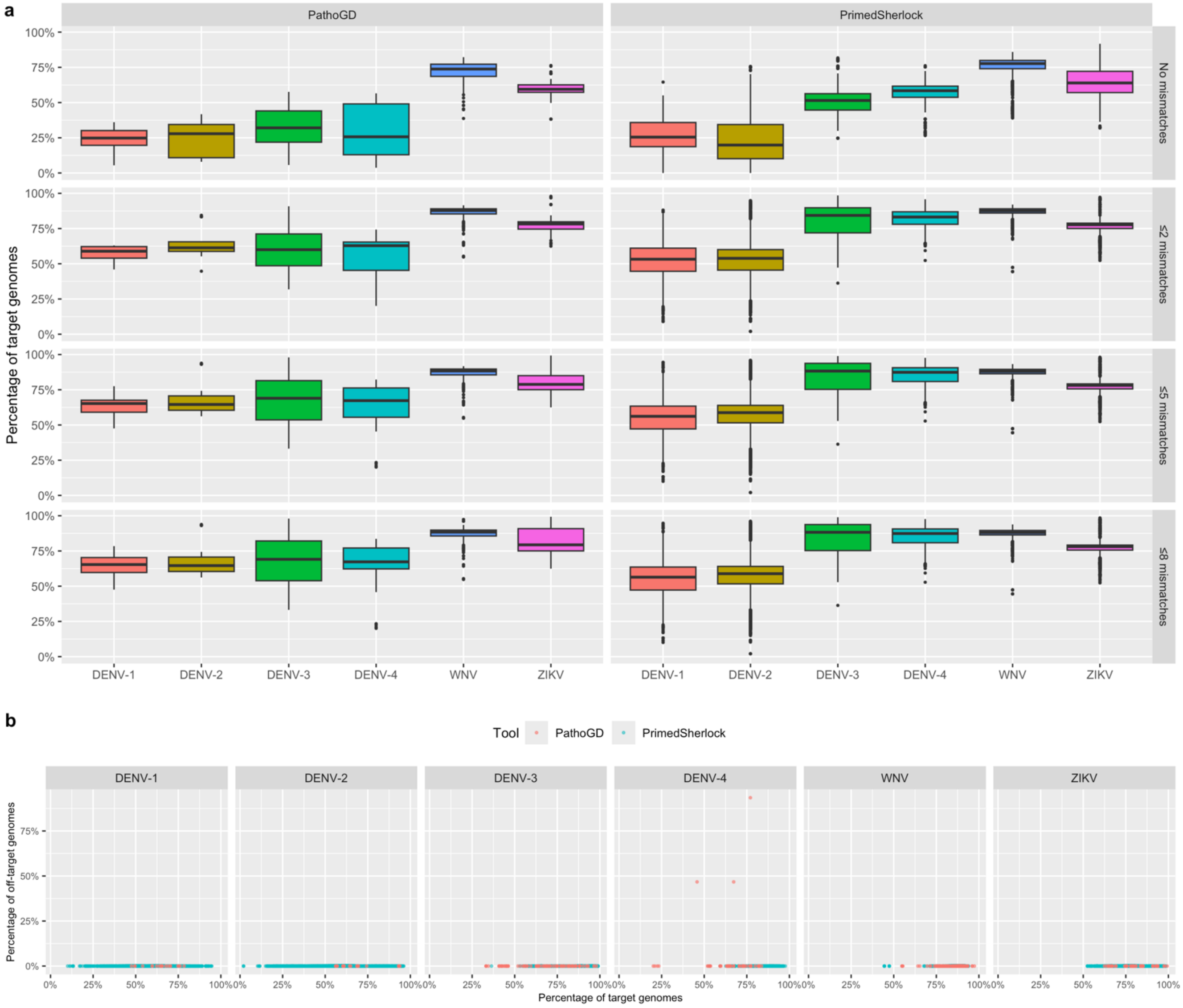


**Figure S6. Comparison of primers designed by PathoGD and PrimedRPA/PrimedSherlock for six viruses. a)** Percentage of target genomes covered by PathoGD and PrimedRPA/PrimedSherlock primers for six viruses at four thresholds of total allowed mismatches across both primers: no mismatch, ≤2 mismatches, ≤5 mismatches and ≤8 mismatches. The centre line of the boxplots corresponds to the median, the lower and upper hinges correspond to the first and third quartiles, respectively, and the ends of the whiskers represent values no further than 1.5x of the interquartile range. **b)** Scatterplot of PathoGD and PrimedSherlock primer specificity. Each point represents the percentage of target and off-target genomes covered by a single primer set, allowing for 8 total mismatches across both primers.

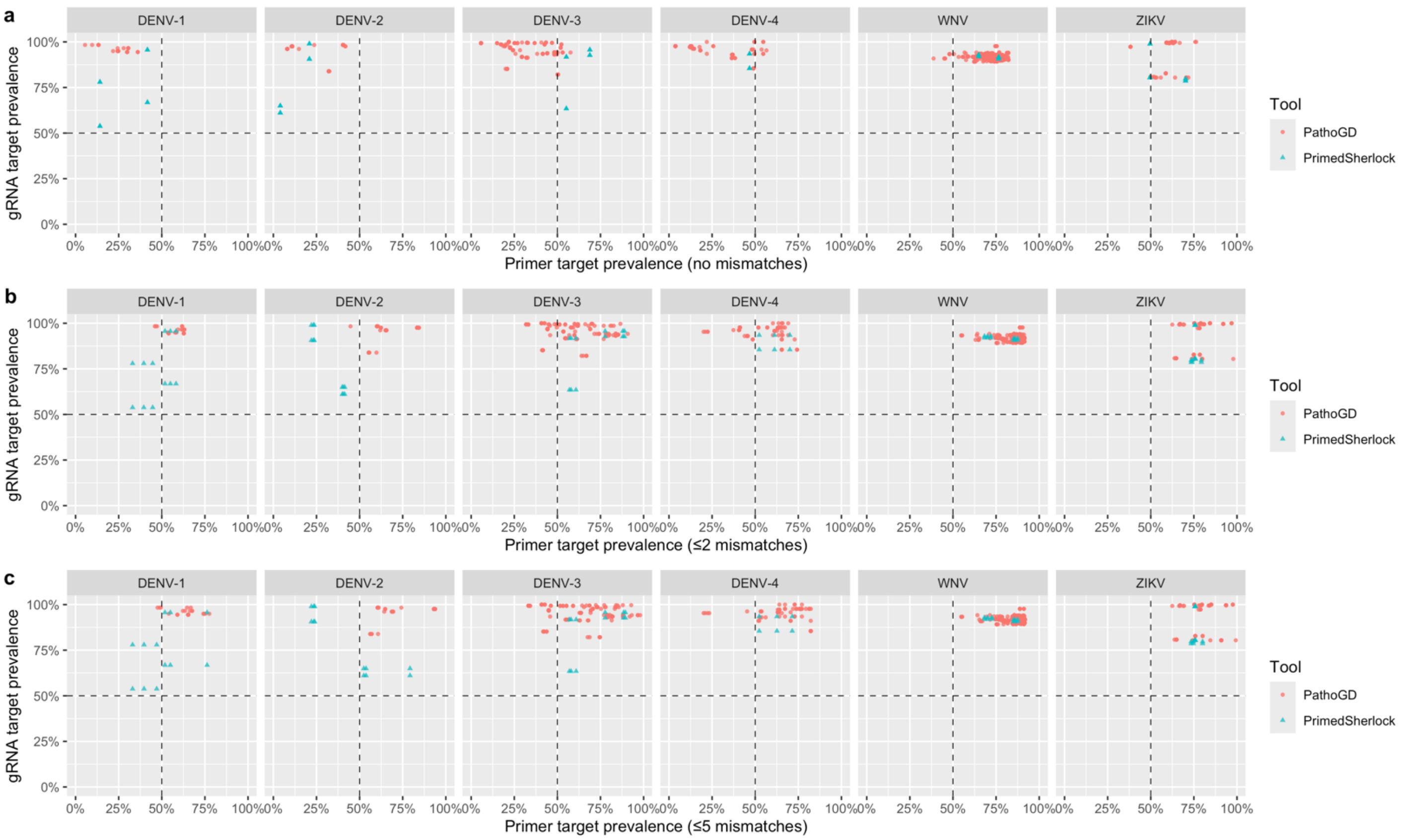


**Figure S7. Prevalence and conservation of primer and gRNA binding sites across target genomes of six viruses for PathoGD and PrimedSherlock designs. a)** The percentage of target genomes covered by each primer/gRNA combination. PrimedSherlock designs were reported as the best primer and gRNA combinations. **b)** and **c)** Similar to (a) but with ≤2 mismatches and ≤5 mismatches allowed across the primer binding sites, respectively. For gRNA target prevalence, a maximum of one mismatch to the target genome was allowed. Black dotted lines denote the 50% cutoff for prevalence of primers and gRNAs.

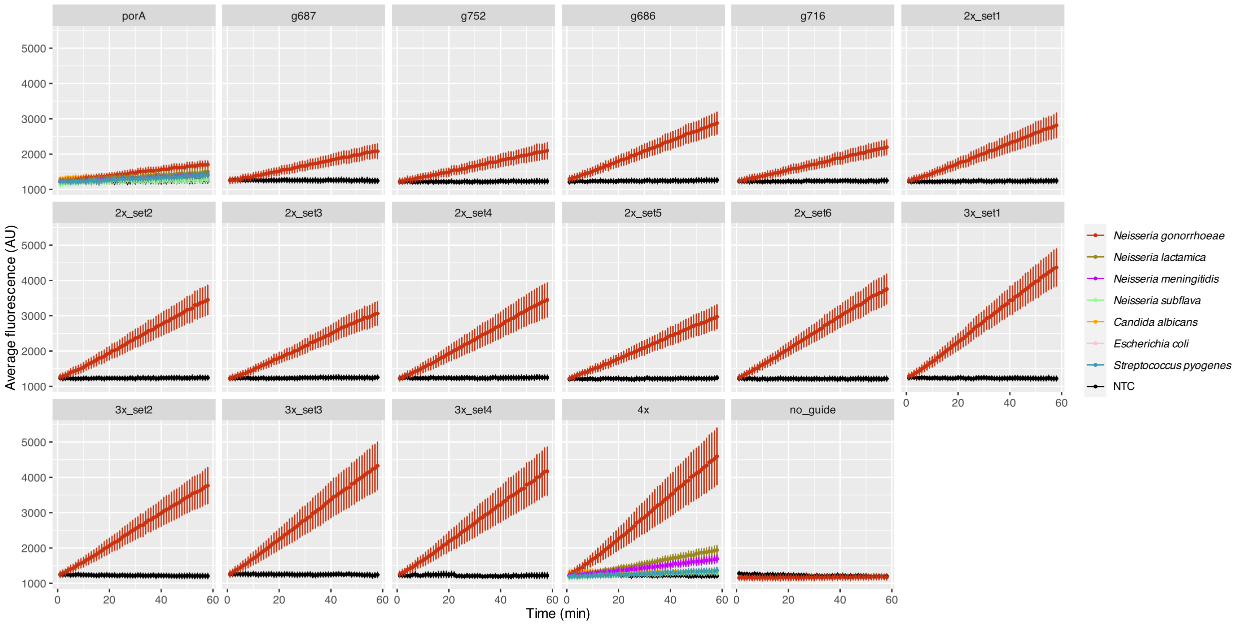


**Figure S8.** Individual fluorescence profiles across 58 min for *porA* and six multi-copy gRNAs, individually and in pooled combinations. Data shown is average fluorescence ± standard error of mean (SEM) across at least three experimental repeats.

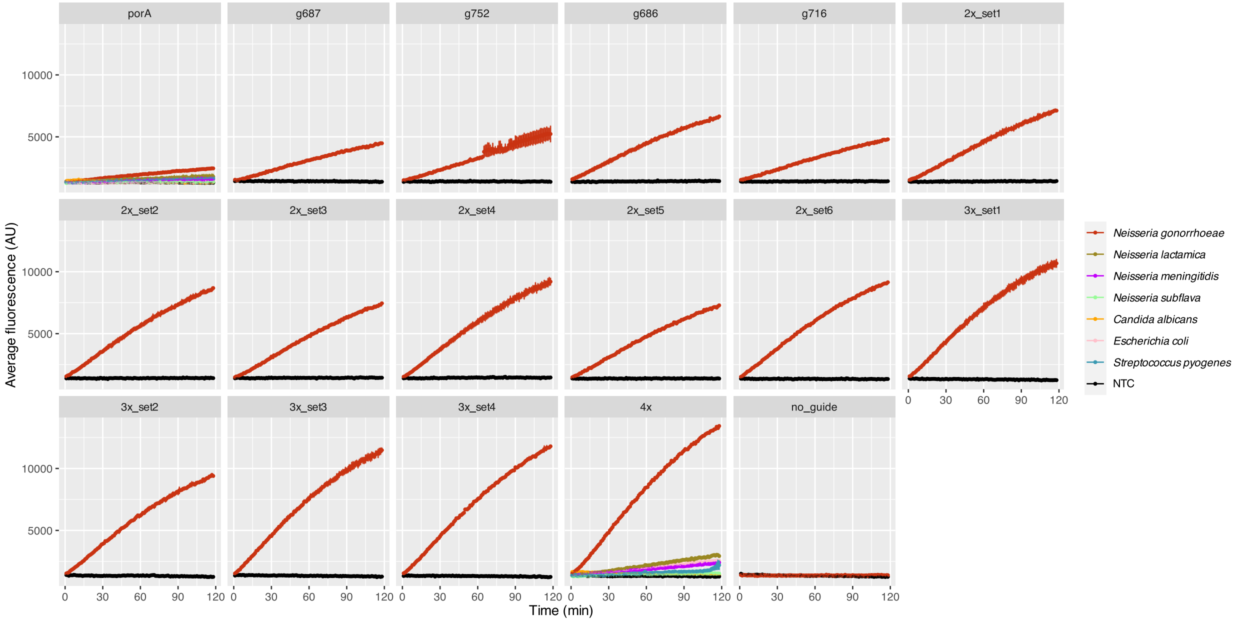


**Figure S9.** Individual fluorescence profiles across 118 min for *porA* and six multi-copy gRNAs, individually and in pooled combinations. Data shown is average fluorescence ± standard error of mean (SEM) of two technical replicates.
